# Predicting drug-target interactions using multi-label learning with community detection method (DTI-MLCD)

**DOI:** 10.1101/2020.05.11.087734

**Authors:** Yanyi Chu, Xiaoqi Shan, Dennis R. Salahub, Yi Xiong, Dong-Qing Wei

## Abstract

Identifying drug-target interactions (DTIs) is an important step for drug discovery and drug repositioning. To reduce heavily experiment cost, booming machine learning has been applied to this field and developed many computational methods, especially binary classification methods. However, there is still much room for improvement in the performance of current methods. Multi-label learning can reduce difficulties faced by binary classification learning with high predictive performance, and has not been explored extensively. The key challenge it faces is the exponential-sized output space, and considering label correlations can help it. Thus, we facilitate the multi-label classification by introducing community detection methods for DTIs prediction, named DTI-MLCD. On the other hand, we updated the gold standard data set proposed in 2008 and still in use today. The proposed DTI-MLCD is performed on the gold standard data set before and after the update, and shows the superiority than other classical machine learning methods and other benchmark proposed methods, which confirms the efficiency of it. The data and code for this study can be found at https://github.com/a96123155/DTI-MLCD.

## 1. Introduction

For drug development, drug discovery (i.e. finding potential new drugs) and drug repositioning (i.e. obtaining old drugs with new efficacy) are two important strategies with heavy cost [2], and an important step to achieve them is predicting DTIs. In recent years, many studies have applied the popular machine learning technology to realize intelligent medical treatment, which has accelerated the process of drug development to a certain extent. For DTIs prediction, the use of machine learning techniques can not only reduce the experimental scope of experimental research but also play a guiding role in experimental research.

There are many review articles [3–7] summarizing the progress of machine learning methods in the field of DTIs prediction in recent years, and the binary classification method is an important branch. For the binary classification method [8–41], drug-target pairs and interactions are treated as samples and labels, respectively. It describes the drug-target pair by encoding drugs and targets as the feature vector, then, predicts DTIs by building a binary classifier. In addition to the binary classification methods, there are network inference methods [42–55], matrix factorization methods [56–63], kernel-based methods [64–68], restricted Boltzmann machine method [69], collaborative filtering method [70], clustering method [71], label propagation method [72], etc. It is worth noting that many of these other methods can be attributed to the binary classification method in a sense. For example, the network inference method regards the DTIs prediction problem as the bipartite network inference problem, and infers missing edges to achieve DTIs prediction. If the missing edges are regarded as negative samples and the existing edges are regarded as positive samples, it is converted into a binary classification problem.

For the binary classification method, it requires the participation of positive and negative samples, so unknown DTIs are often treated as negative samples. This negative sample construction strategy will not only introduce noise but also cause data imbalance as a large number of negative samples. Besides, it is also faced with excessive computational load and overfitting due to the redundant feature space and extremely high feature dimensions. For example, 10 drugs and 10 targets will be combined into 10 × 10 = 100 samples, and the same drug or target in different samples has the same feature vector, that is, the feature vector of each drug or target will appear 10 times in the feature space of 100 samples. To reduce the above difficulties, the application of multi-label learning to DTI prediction problems is worth exploring. The multi-label classification problem trains a model that maps the input feature vector to more than one label. Transform the above binary classification example into a multi-label classification problem, described as: 10 drugs (or targets) as samples, and 10 targets (or drugs) as labels. The input feature is only a description of 10 drugs (or targets). Then use the multi-label learning algorithm to predict drug targets (or drugs that can interact with the specific target). Obviously, unlike the binary classification problem, it only requires information about the drug (or target) to predict a series of DTIs. The experiments in this study prove that its performance is very competitive with the binary classification problem, and its speed is much higher than that of the binary classification method, especially for large data sets. Until now, there are few applications and a lot of space for exploring multi-label learning applied in the DTIs prediction problem. DrugE-Rank [73] is a method using the “Learning To Rank” paradigm to model the DTIs prediction problem as a multi-label task. A study [74] uses multi-task deep neural networks for drug targets prediction, and firstly uses extended connectivity fingerprints with radius 12 as drug representation. Moreover, to overcome the training difficulties caused by too many labels in multi-label learning, Pliakos et al. [75] proposed three multi-label learning methods for DTI prediction, which use k-means for label division.

On the other hand, the gold standard data set currently used in the field of DTIs prediction is the data set collected by Yamanishi in 2008 [76], named Yamnishi_08. After 12 years, a large number of new DTIs have been discovered, but they were not considered. As we all know, positive samples (i.e. DTIs) are essential for method construction. The missing of positive samples not only introduces error in the modeling process, but also hides a great risk of false negatives during the model evaluation, making the unknown bias between research results and the actual results. For this point, Keum and Nam [11] updated these datasets among the original drugs and targets. However, in reality, it cannot be limited to the original drugs and targets, and the DTI between new drugs and targets should also be considered.

This study updates the gold standard data set of drugs, targets, and DTIs. In addition, we proposed the multi-label learning with community detection method for DTIs prediction (DTI-MLCD) and tested it on four original and updated gold standard data sets. The proposed DTI-MLCD first uses the community detection algorithm to divide the target space into multiple subspaces, then applies multi-label learning on each subspace, and finally performs DTIs prediction. Comparison with traditional machine learning methods and other benchmark DTIs prediction methods confirms the effectiveness of the proposed DTI-MLCD method.

## 2. Material and Methods

### 2.1. Problem description

This study divides the DTIs prediction problem into two sub-tasks: (a) drug discovery, which predicts new drugs, named T_D_; (b) drug repositioning, which predicts new targets, named T_T_. These two tasks are regarded as multi-label classification problems, described below.

For task T_D_, suppose *X*_*D*_ =R^*d*^ and *Y*_*T*_ = {*y*_1_, *y*_2_,…, *y*_*p*_} denote the *d*-dimensional drug instance space and the label space with *p* possible target class labels. This task is to learn a function 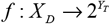 from the multi-label training set *D* = {(*x*_*D*,*i*_, *y*_*T*,*i*_) |1 ≤ *i* ≤ *m*}, where *m* is the number of samples. For each sample (*x*_*D*,*i*_, *y*_*T*,*i*_), *x*_*D*,*i*_ ∈ *X*_*D*_, it is a *d*-dimensional feature vector and *y*_*T*,*i*_ ∈ *Y*_*T*_ is the label set associated with *x*_*D*,*i*_. For drug instances of the test set, the multi-label classifier *f* (·) predicts the proper labels. The task T_T_ can be defined by analogy.

### 2.2 Data sets

Yamnishi_08 derives from the KEGG BRITE [77], BRENDA [78], SuperTarget [79], and DrugBank [80] databases. It consists of four DTI datasets. These datasets are different according to the protein targets, namely nuclear receptor (NR), G-protein-coupled receptor (GPCR), ion channel (IC), and enzyme (E). To update these datasets, we collect new drugs, new targets, and new DTIs using KEGG BRITE, UniProt [81], and DrugBank databases in this study. The steps are divided into two parts: data integration and data cleaning. Data integration is achieved through web crawler technology. First, the DTI data corresponding to the 4 types of targets is obtained from the KEGG BRITE database and merged with Yamanishi_08 to prevent the loss of information in the SuperTarget and BRENDA databases. Then, use the UniProt database as the connection database of KEGG BRITE and DrugBank, search the DrugBank database for each target obtained in the previous step, and add drugs and corresponding DTIs that are not in KEGG BRITE and Yamanishi_08. Next, search all known drugs one by one to maximize the DTI integrity of existing drugs and targets. After obtaining the integrated data, we deleted useless, invalid, and redundant data, including non-small molecule drugs (such as biotechnology drugs), mixed drugs, drugs with the same or unknown structure, and drugs with unknown end groups in the structure. It is worth noting that all drugs in the updated data set are approved drugs. The code for updating the dataset has been published on the GitHub page. Some statistics of the original gold standard and new updating four data sets are shown in Table 1.

**Table 1.** Statistics of the original and updating four data sets. The NR is short for the nuclear receptor, GPCR for the G-protein-coupled receptor, IC for the ion channel, and E for the enzyme. Besides, the n represents the amount, D represents degree, and the subscripts d and t represent drug and target, respectively.

| Data sets | | $n_d$ | $n_t$ | $n_{interaction}$ | Density (%) | $D_d$ | $D_t$ | $D_d=1$ (%) | $D_t=1$ (%) |
| --- | --- | --- | --- | --- | --- | --- | --- | --- | --- |
| NR | Original | 54 | 26 | 90 | 6.41 | 1.67 | 3.46 | 72.22 | 30.77 |
|  | Updated | 541 | 33 | 886 | 4.96 | 1.64 | 26.85 | 65.99 | 18.18 |
| GPCR | Original | 223 | 95 | 635 | 3.00 | 2.85 | 6.68 | 47.53 | 35.79 |
|  | Updated | 1680 | 156 | 5383 | 2.05 | 3.20 | 34.51 | 46.13 | 14.74 |
| IC | Original | 210 | 204 | 1476 | 3.45 | 7.03 | 7.24 | 38.57 | 11.27 |
|  | Updated | 765 | 238 | 6385 | 3.51 | 8.35 | 26.83 | 21.70 | 8.82 |
| E | Original | 445 | 664 | 2926 | 0.99 | 6.58 | 4.41 | 39.78 | 43.37 |
|  | Updated | 1777 | 1411 | 7371 | 0.29 | 4.15 | 5.22 | 45.24 | 37.99 |

### 2.3. Features

#### 2.3.1. Drug representation

Many representations can be used to describe drugs, in general, these can be categorized into two types: molecular descriptors (MDs), molecular fingerprints (MFs). To explore the drug representation that is most suitable for this study, we used some open source tools commonly used in DTI prediction to generate MDs and MFs. For the MDs or MFs generated by different software, this study treats them as different drug representations. The tools used in this study are CDK [82], Pybel [83], RDKit [84], and PaDEL [85]. The MDs generated by the above tools are called MD_CDK, MD_PYB, MD_RDK, and MD_PAD. Their dimensions are 275, 24, 196, and 1875, respectively. Further, we combine these four types of MDs as a new type of MDs, called MD_MER. Currently, MFs are always divided into three categories [86]: (a) topological path-based fingerprint. The representative FP2 [87] (MF_FP2) used in this study; (b) topological circular fingerprint. ECFP4 [88] (MF_EC4) and ECFP8 [88] (MF_EC8) are used as their representativeness; (III) substructure key-based fingerprint. MACCS [89] (MF_MAC) and PubChem fingerprint [90] (MF_PCP) are used as their popularity in DTIs prediction. The dimension of them is 1024, 2048, 2048, 167, and 881, respectively. In addition to the MDs and MF, we also used the Word2vec-inspired feature [33] (W2V), which extracts word information from drug SMILES.

Further, we perform the feature combination of the above three types of features, because the complementarity between these three types of features may help enhance performance. In this process, we fuse the feature selection to obtain clean, highly complementary, and less redundant combined features.

#### 2.3.2. Target representation

This study uses three types of target sequence-derived representations commonly used in DTIs prediction studies. The first is Composition, Transition, and Distribution (CTD), which is represented as the 504-dimensional feature vector obtained by PROFEAT web server [91]. The second is 1437 default protein descriptors generated by PROFEAT, named PRO. Besides CTD, it also includes amino acid composition, dipeptide composition, autocorrelation, quasi-sequence-order, amphiphilic pseudo-amino acid composition, and total amino acid properties. The third is the protein domain fingerprint (PDF), which is extracted from the PFAM v31.0 database [92]. For different data sets, we extracted different numbers of domains. The feature vector dimension of targets in NR, GPCR, IC, and E is 30, 61, 1404, and 2182, respectively. In addition, the feature combination is also performed.

### 2.4 Methods

The traditional supervised learning can be regarded as a degenerated version of multi-label learning as each sample is confined to have only one single label. However, the generality of multi-label learning makes harder to design the algorithm. The exponential-sized output space is the core issue of learning, i.e. there are 2^*m*^ possible label sets for *m* labels. And exploiting label correlations or executing label space partition can help it. For this purpose, this study applies the community detection method from social networks to divide label space. Next, each divided label subspace corresponds to a multi-label learning sub-problem, and multiple Label Powerset (LP) multi-label classifiers are jointed to cover the entire label space. The base learner applied in LP is random forest (RF) because of its simplicity, parallelism, and superior capabilities, etc. In this section, we will introduce the typical algorithms of multi-label learning and community detection. The execution steps of the proposed DTI-MLCD method are shown in Figure 1.

**Figure 1.**
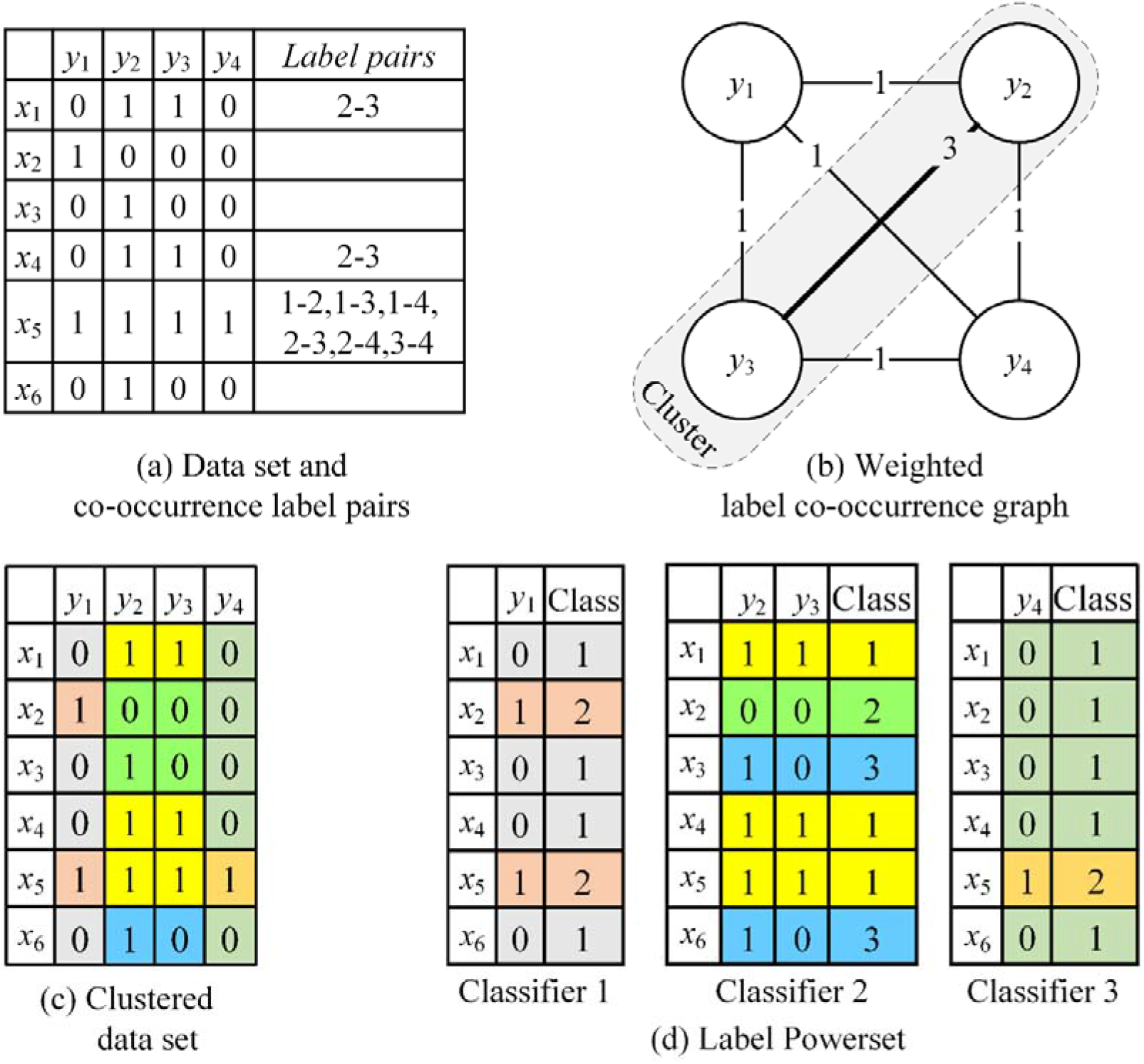
The execution steps of the proposed DTI-MLCD method.

#### 2.4.1. Algorithms of multi-label learning

The multi-label learning algorithm development is the key challenge in multi-label learning researches, although the algorithm has boomed in a big explosion in the past 10 years. A simple categorization is adopted as follows.

The first category is the algorithm adaptation method, which works by fitting the existing algorithm to data and directly tackle the multi-label data. The representative algorithm is Multi-Label *k*-Nearest Neighbor (ML*k*NN) [93]. ML*k*NN is a lazy learning method based on the traditional *k*-Nearest Neighbor. It is now widely used in multi-label classification prediction tasks and has achieved satisfactory results [94, 95]. The second category is the problem transformation method, which works by fitting data to the well-established algorithm and transforming multi-label learning problems into the other learning technique. Binary Relevance (BR) [96], Classifier Chains (CC) [97] and Label Powerset (LP) [98] are representative algorithms in this category. BR tackles multi-label learning problem into multiple independent binary classification problems, where one binary classifier corresponds to one label. It is based on the assumption that labels are independent of each other, and each classifier of this method only recognizes the characteristics related to one label, but cannot identify the characteristics related to all labels. Thus, it is not valid in many fields in reality, which is also the limitation of the BR. CC is proposed based on BR and considers label correlation. It converts the multi-label learning problem into a chain of binary classification problems. The main idea is to add the labels of all previous classifiers to the feature vector of the next training set and pass them to the next classifier. Obviously, the order of labels has a great influence on the prediction result. However, the order of the classifiers in the classifier chain is always random. Unlike BR and CC, LP transforms the multi-label learning task into the multi-class or single-label classification task. In other words, LP models the joint distribution of labels. It treats each label subset in the multi-label training set as a class of a multi-class task, and the prediction will be one of these subsets. Although LP is simple, it has two impractical points that tend to cause over-fitting. One is incompleteness. It can only predict label sets appearing in the training set, and is powerless for other label sets. The other is inefficiency. As the number of labels increases, it may face high complexity because of the increase in the number of label subsets, and the high imbalance of samples in each class or subset.

To overcome the shortcoming of LP while retaining its simplicity, the idea that dividing the label space into multiple subspaces and applying the LP algorithm in these subspaces has been proposed [99], which can be seen as combining ensemble learning with LP. This is the design principle of random *k*-labelsets (RA*k*EL) [99]. RA*k*EL divides the overall label set into multiple size-*k* label subsets randomly and implements LP on each label subspace to ensure computational efficiency. Then, it ensembles several LP classifiers to guarantee the completeness of the prediction. However, an obvious disadvantage of RA*k*EL is the random partition strategy, which makes the label correlation controlled only by *k*, and not considered training data.

To consider the correlation among labels informatively, the data-driven clustering algorithm is used instead of the random partition strategy. Moreover, it has been confirmed that the data-driven method is superior to random selection for the label space division in multi-label classification problems [100]. Especially, the community detection method has been well applied to multiple benchmark data sets for multi-label learning, it divides the label space in a data-driven manner [100]. Thus, this study discusses the application of five classic community detection algorithms in DTIs prediction.

#### 2.4.2. Execution of community detection

Community detection is to find tightly connected community structures in complex network structures, that is, to discover clusters of nodes in the network [100]. In this study, the goal of using the community detection method is to divide label space with a data-driven approach. For this purpose, the community detection method is used based on the weighted co-occurrence graph which is constructed based on training data.

##### 2.4.2.1. Construct the weighted label co-occurrence graph

Defining the weighted undirected co-occurrence graph, where vertices represent the label set, edges represent label pairs that occur together at least once in the training label set, and the weight assigned to each edge is defined as the number of samples that have both labels. The visualization of the weighted label co-occurrence graph is shown in Figure 1 (a) and (b).

##### 2.4.2.2. Algorithms of community detection

There are many algorithms for community detection. This study uses only five typical algorithms.

The modularity-based approach is a very important branch of the community detection algorithm. It works through maximizing the modularity to implement label space division. Modularity [101] is a measure to describe the quality of the community partition. A relatively good partition has a higher degree of similarity in the community. However, finding the maximum of modularity is NP-hard [102], so we employ three approximation-based techniques instead.

*The fast greedy algorithm (FGA)* [103] is based on the greedy algorithm to search the maximization of modularity, which iteratively merges communities from a single instance. With each iteration, this method merges two communities to achieve the greatest contribution to modularity. When the modularity value of the current community can no longer increase as the community merges, it is defined as convergence.

The *multi-level algorithm (MLA)* [104] is a bottom-up algorithm. In the beginning, each vertex is a separate community, and the vertices move iteratively between the communities by maximizing the local contribution of the vertices to the overall modularity. When modularity is not increased by any movement, each community in the original graph shrinks to a vertex while maintaining the total weight of adjacent edges, and then the process enters the next level. When communities shrink to vertices and the modularity can no longer be increased, the algorithm will stop.

In addition to the modularity-based algorithm, we also use three other algorithms concerning flow.

The *label propagation algorithm (LPA)* [105] is based on the graph semi-supervised learning algorithm, which simulates the diffusion of flow on the network through the diffusion of labels. In the graph, each vertex is assigned a unique label. Next, the tag of every vertex is updated iteratively with the majority tag assigned to the neighbors of the elements. The update order for each iteration is random. The convergent criterion of the algorithm is when all vertex tags are consistent with the most frequent tags in their neighborhood.

The *walk trap algorithm (WTA)* [106] is a bottom-up approach based on random walks. One intuition is that when performing random walks on the graph, it is easy to fall into the dense connection of the graph, which can be regarded as a community. Consider each node as a community, and then calculate the random walk distance or flow distance between all communities with connected edges. Then, take two communities that are connected and have the shortest random walk distance to merge, recalculate the distance between the communities, and then iterate until all nodes are put into the same community.

The *infomap algorithm (IMA)* [107] believes that a good community division should make the average description length of the flow the shortest. It divides the graph by calculating the minimum value of the map equation, where the map equation corresponds to the length of the information description corresponding to the partition.

### 2.5. Performance evaluation

The performance evaluation metrics of multi-label learning are much complex than binary classification [108]. Following the previous researches, this study adopts AUC and AUPR as performance evaluation metrics. It is convenient for comparison with other methods. It is worthy to note that the AUPR is more reliable metrics as a severe punishment on false positive instances for high imbalanced data. Therefore, the discussion in this article focuses on AUPR.

### 2.6. Stratified cross-validation (SCV)

Cross-validation is a typical method to do model selection. For multi-label data, many labels have class imbalance characteristics [109] that each data set has a large number of label sets, and most label sets only contain a small number of samples (Table 2). In this case, the random partitioning strategy used in standard cross-validation may result in some labels having no positive samples in a divided subset. Such a subset will not only affect the accuracy of the model, but may also cause the computational error.

**Table 2.**
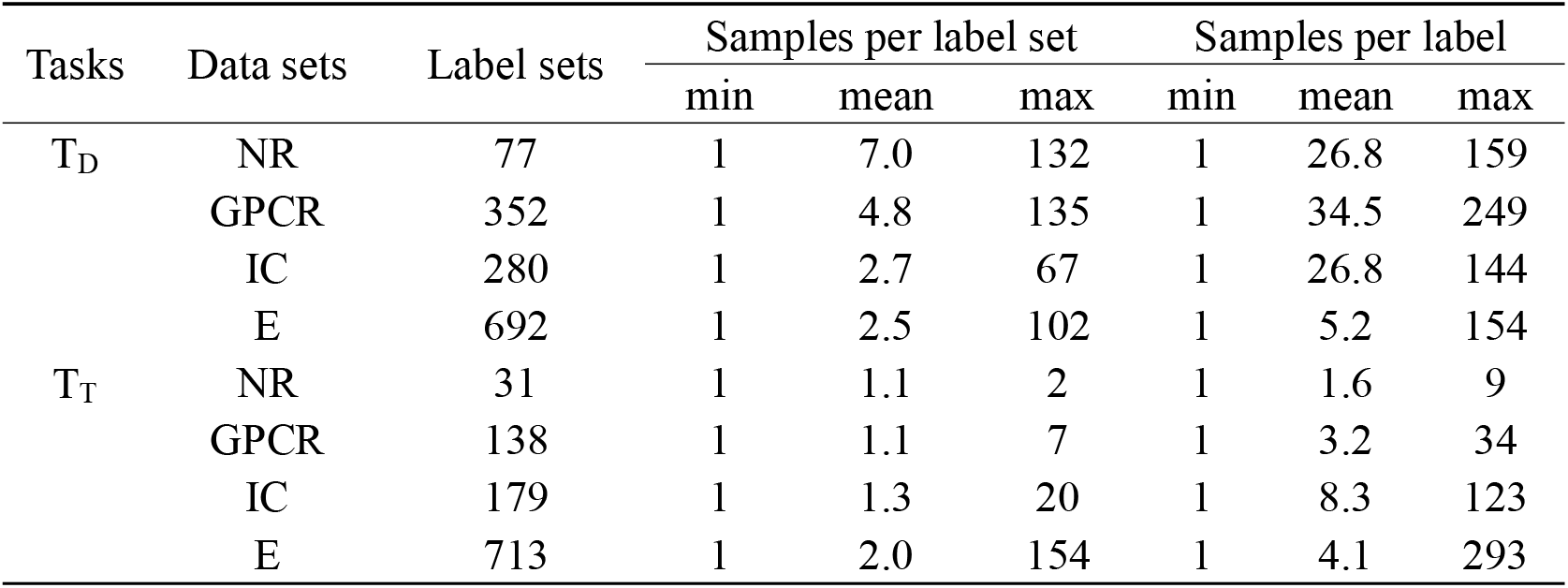
Statistics for labels of four multi-label data sets. The data in the table is the number of corresponding row and column headings. For the Data sets column, the NR is short for nuclear receptor, GPCR for G-protein-coupled receptor, IC for ion channel, and E for enzyme. For the Tasks column, the T_D_ is predicting new drugs, T_T_ is predicting new targets.

To overcome the above dilemma, a stratified sampling strategy in cross-validation is a proven solution [109, 110], called stratified cross-validation (SCV). Furthermore, the 10-fold SCV has been proved the best method in model selection from the perspective of statistical inference [110]. To ensure the confidence of the results, we performed 5 simulations on 10-fold SCV using different random seeds.

### 2.7. Hypothesis test

When comparing multiple algorithms on a set of data sets, Demšar [111] recommends using the non-parametric Friedman rank test [112, 113] which based on algorithm ranking. However, the Friedman rank test can only tell us whether there is a significant difference among algorithms, but cannot specify which algorithms have performance differences. Therefore, post-hoc analysis is needed to locate specific algorithms with differences. For the Friedman rank test, the commonly used post-hoc test method is the Nemenyi test [114], named Friedman-Nemenyi test. This method can indicate whether there is a significant difference between the two algorithms based on the significance level α.

## 3. Results and Discussion

### 3.1. Selecting drug representation

We guess that for different data sets, the most suitable drug representation method is different. So far no other articles have explored this, and our following experiments prove this conjecture. And this phenomenon makes us apply different feature representation methods in different data sets.

To achieve this goal, the experiment is conducted on the basic learning algorithm LP for each updated data set, and the same parameter settings were used. The AUPR and AUC are shown in Figure 2. However, AUPR is the focus as its more reliable, and its lower value is more valuable than high AUC for discussion and comparison.

**Figure 2.**
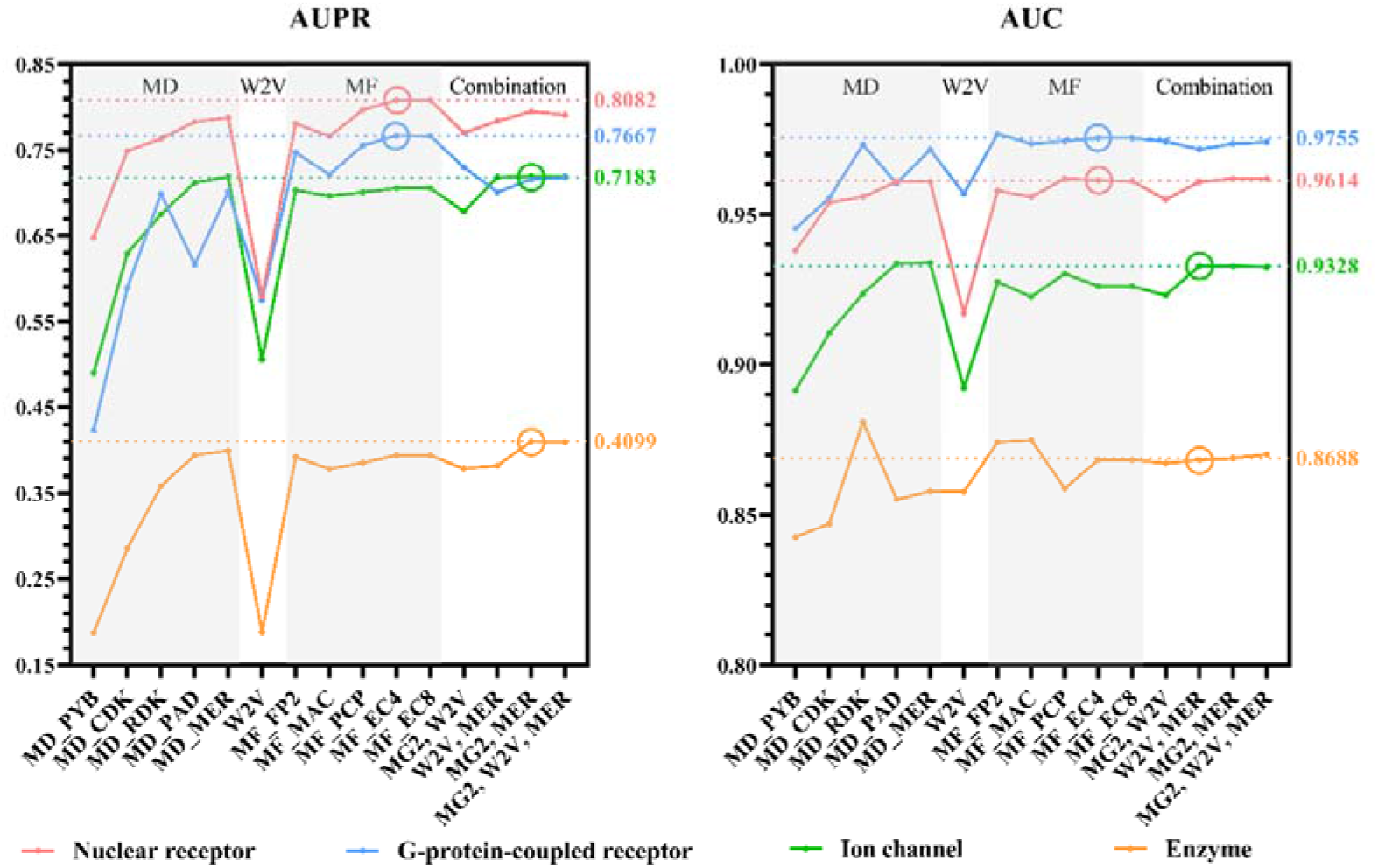
The performance among different drug representations.

For MDs, on the four data sets, as the dimension of drug representation increases, the prediction performance tends to be higher because of the more information it describes. For MFs, MF_EC4 is the best MF among all four data sets, and it has been proved that is sufficient to describe chemical molecules [115]. Further, the result reveals the topological circular fingerprint is better than the other two categories in this study. Next, the feature combination procedure has been performed. There are 4 combinations of MD_MER, MF_EC4, and W2V. Figure 4 indicates that the performance of any drug representation after adding W2V was lower than that without W2V.

For different data sets, this study selects the drug representation with the best AUPR as the feature vector. NR and GPCR use MF_EC4, IC and E use the combination of MF_EC4 and MD_MER.

### 3.2. Selecting target representation

We have adopted the same strategy as drugs, that is, there is no best target representation method, only the most suitable feature representation in a specific situation. Therefore, we also compare target representation methods under four updated data sets and select the most suitable features for each data set according to AUPR.

According to Figure 3, it is obvious that the performance of CTD and PRO is close, probably because both of them are generated by the PROFEAT web server, and CTD is a subset of PRO. Further, for the combination of CTD or PRO with PDF, the performance is also close. Besides, on the NR and GPCR data sets, PDF appears to be a significant trough. Because the protein domain information is too little to fully describe the target. Also, its lower dimension than CTD and PRO makes it have little effect on the performance of feature combinations. On the contrary, on the IC and E data sets, the performance of PDF is significantly improved compared to CTD and PRO as its rich protein domain information. Therefore, PDF dominates the performance of feature combinations.

**Figure 3.**
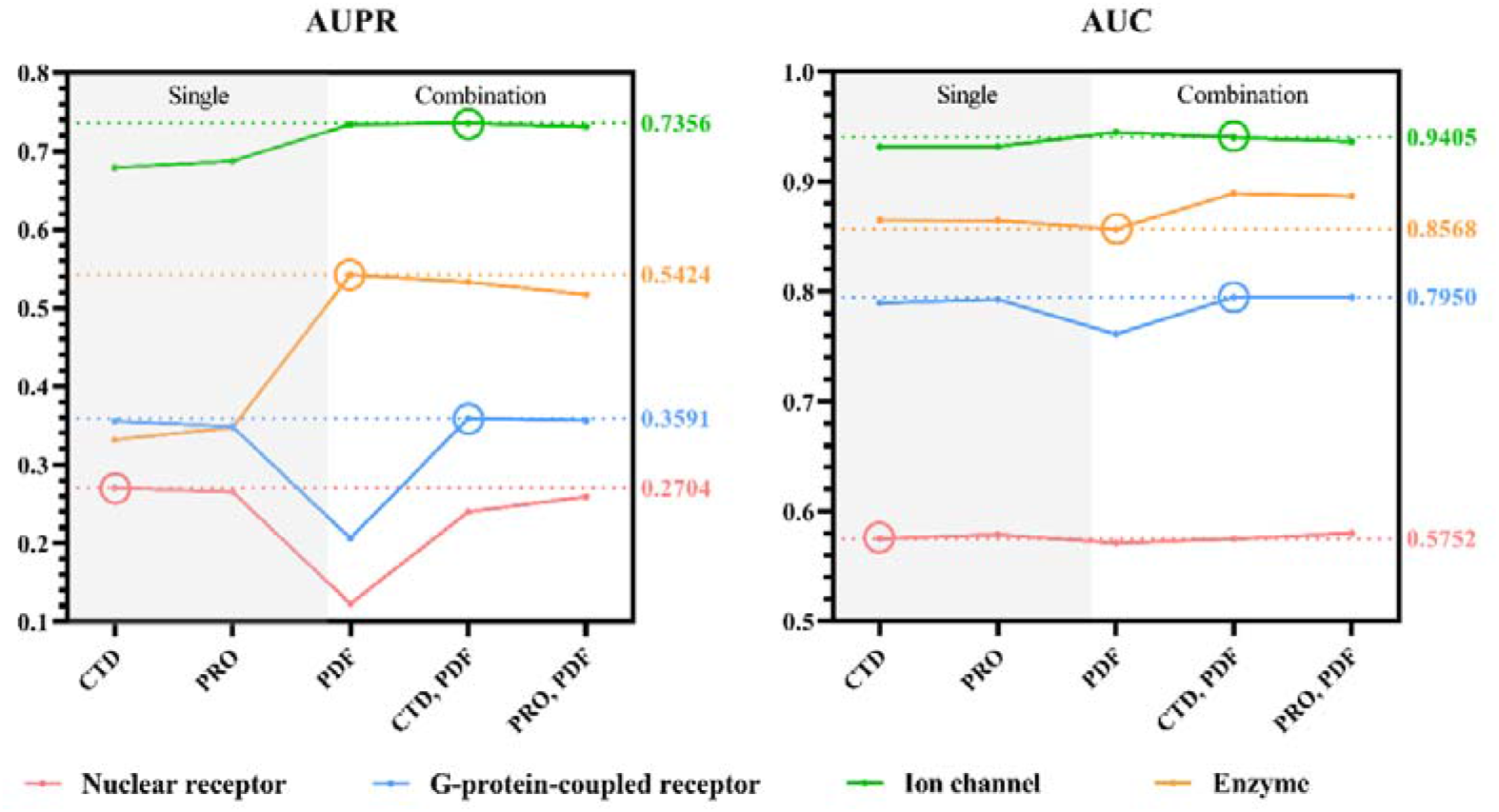
The performance among different target representations.

Finally, we chose the most suitable target representation method for each data set. For NR, the most suitable target representation method is CTD, for IC is PDF, for GPCR and E is the combination of CTD and PDF.

### 3.3. The DTI-MLCD and classical machine learning methods in updated data sets

This study proposed the DTI-MLCD method which applies five data-driven community detection algorithms as label partitioning methods and ensembles them into the multi-label learning method. We explain the superiority of DTI-MLCD from two aspects. The first is the comparison of label partitioning algorithms. For data-driven label partitioning method, *k*-means is always used due to its simplicity and popularity, and has been applied with *k* ∈{2, 4,8,16, 32} to solve the DTIs prediction problem [75]. So we use *k*-means as the benchmark label partition method to compare with community detection algorithms. To be more convincing, we expanded the value range of *k* from 2 to the number of the label set, and used the silhouette coefficient score as a measure of label division quality. The *k* value that maximizes the silhouette coefficient score will be used as the optimal number of clusters. The other is to compare the DTI-MLCD algorithm with other classic machine learning algorithms, (a) multi-label algorithms: ML*k*NN, BR, CC, LP, and RA*k*EL, and (b) binary classification algorithms: RF, extremely randomized Trees (ERT), and Gaussian naïve Bayes (GNB).

The results of the above methods on the four updated data sets are listed in Table 3 (task T_D_) and Table 4 (task T_T_), respectively. In the results, the proposed DTI-MLCD is superior to other machine learning methods in most cases. The reason why LP performs better than DTI-MLCD on the NR data set under the T_T_ task is that NR has few label sets, and both each label set and each label has very few samples (see Table 2 for details), but it has 541 labels. Therefore, only a single LP algorithm can achieve better results, but after adding the community detection algorithm, it will cause overfitting. On the other hand, although the binary classification methods RF and ERT have achieved competitive results with DTI-MLCD, however, this research experiment proves that its long calculation time will make it difficult to achieve optimal performance through fine-tuning. Further, the Friedman-Nemenyi test with a significance level of 0.05 confirmed the significant differences among methods. All the five proposed methods are at the forefront of the ranking, and the overall performance of FGA is slightly lower than the other four proposed methods.

**Table 3.** The results of the proposed methods and other classical machine learning methods for task T_D_ (i.e. predicting new drugs). WTA, IMA, LPA, MLA, and FGA are community detection algorithms in the proposed DTI-MLCD method. *k*-means is the benchmark method that can replace community detections. BR, CC, LP, RA*k*EL, ML*k*NN, RF, ET, and GNB are baseline methods.

| Algorithm | AUC |  |  |  | AUPR |  |  |  |
| --- | --- | --- | --- | --- | --- | --- | --- | --- |
|  | NR | GPCR | IC | E | NR | GPCR | IC | E |
| FGA | 0.9613 | 0.9738 | 0.9349 | 0.8840 | **0.8135** | 0.7721 | 0.7184 | 0.4148 |
| IMA | 0.9611 | 0.9766 | 0.9358 | 0.8768 | 0.8129 | **0.7765** | **0.7194** | 0.4165 |
| LPA | 0.9611 | 0.9763 | 0.9345 | 0.8833 | **0.8135** | 0.7755 | 0.7179 | 0.4173 |
| MLA | 0.9614 | 0.9745 | 0.9347 | 0.8833 | 0.8134 | 0.7734 | 0.7186 | 0.4165 |
| WTA | 0.9611 | 0.9744 | 0.9355 | 0.8839 | 0.8129 | 0.7722 | 0.7187 | **0.4184** |
| $k$-means | 0.9629 | 0.9754 | 0.9352 | 0.8771 | 0.8128 | 0.7731 | 0.7178 | 0.4040 |
| BR | 0.9622 | 0.9814 | 0.9372 | 0.8771 | 0.8115 | 0.7307 | 0.6914 | 0.4040 |
| CC | 0.9610 | 0.9767 | 0.9346 | 0.8664 | 0.8109 | 0.7219 | 0.6845 | 0.3822 |
| LP | 0.9614 | 0.9755 | 0.9328 | 0.8688 | 0.8082 | 0.7667 | 0.7193 | 0.4099 |
| RAkEL | 0.9532 | 0.9735 | 0.9306 | 0.8736 | 0.8004 | 0.7724 | 0.7048 | 0.4034 |
| MLkNN | 0.9363 | 0.9575 | 0.8356 | 0.7962 | 0.6699 | 0.6340 | 0.1644 | 0.0454 |
| RF | 0.9626 | 0.9754 | 0.9423 | 0.8983 | 0.8102 | 0.7730 | 0.7113 | 0.3238 |
| ERT | 0.9616 | 0.9688 | 0.9314 | 0.8786 | 0.8102 | 0.7571 | 0.7049 | 0.3546 |
| GNB | 0.6818 | 0.7037 | 0.5015 | 0.5273 | 0.3732 | 0.3730 | 0.4197 | 0.0054 |

**Table 4.** The results of the proposed methods and other classical machine learning methods for task T_T_ (i.e. predicting new targets). WTA, IMA, LPA, MLA, and FGA are community detection algorithms in the proposed DTI-MLCD method. *k*-means is the benchmark method that can replace community detections. BR, CC, LP, RA*k*EL, ML*k*NN, RF, ET, and GNB are baseline methods.

| Algorithm | AUC |  |  |  | AUPR |  |  |  |
| --- | --- | --- | --- | --- | --- | --- | --- | --- |
|  | NR | GPCR | IC | E | NR | GPCR | IC | E |
| FGA | 0.5715 | 0.8027 | 0.9489 | 0.8593 | 0.2311 | 0.3702 | 0.7468 | 0.5669 |
| IMA | 0.5748 | 0.8002 | 0.9476 | 0.8598 | 0.2409 | 0.3683 | **0.7663** | 0.5669 |
| LPA | 0.5759 | 0.8048 | 0.9459 | 0.8591 | 0.2494 | **0.3785** | 0.7518 | 0.5670 |
| MLA | 0.5657 | 0.8062 | 0.9478 | 0.8640 | 0.2177 | 0.3759 | 0.7609 | **0.5677** |
| WTA | 0.5745 | 0.8002 | 0.9463 | 0.8642 | 0.2401 | 0.3746 | 0.7574 | 0.5673 |
| $k$-means | 0.5611 | 0.7893 | 0.9382 | 0.8639 | 0.2383 | 0.3693 | 0.7174 | 0.5668 |
| BR | 0.5617 | 0.7892 | 0.9382 | 0.8639 | 0.2352 | 0.3694 | 0.7174 | 0.5673 |
| CC | 0.5647 | 0.7580 | 0.9183 | 0.8563 | 0.2360 | 0.2424 | 0.6475 | 0.5152 |
| LP | 0.5752 | 0.7927 | 0.9403 | 0.8568 | **0.2704** | 0.3670 | 0.7429 | 0.5651 |
| RAKEL | 0.5642 | 0.7902 | 0.9395 | 0.8640 | 0.2352 | 0.3714 | 0.7242 | 0.5670 |
| MLkNN | 0.5470 | 0.7351 | 0.9094 | 0.8053 | 0.1811 | 0.2751 | 0.6414 | 0.3112 |
| RF | 0.6764 | 0.7610 | 0.9511 | 0.8775 | 0.2445 | 0.3104 | 0.7419 | 0.5652 |
| ERT | 0.5804 | 0.7179 | 0.9459 | 0.8404 | 0.2632 | 0.3410 | 0.7650 | 0.5462 |
| GNB | 0.4451 | 0.6566 | 0.5006 | 0.5347 | 0.3149 | 0.3770 | 0.3107 | 0.0035 |

To illustrate the biological explanation of the proposed methods, Figure 4 visualizes the results of six data-driven label partitioning methods that applied to the NR dataset. Although the community structures obtained by different community detection algorithms have their own characteristics, they also have certain similarities. FGA, LPA, and MLA divide 33 labels into 6 communities. Especially, the community structure of FGA and MLA is the same, noted that both FGA and MLA belong to the modularity-based algorithm. In addition, for the random walk-based algorithm, the number of communities obtained by WTA and IMA is relatively large. Moreover, the *k*-means obtains only 4 communities, and the community structure is very different from community detection algorithms.

**Figure 4.**
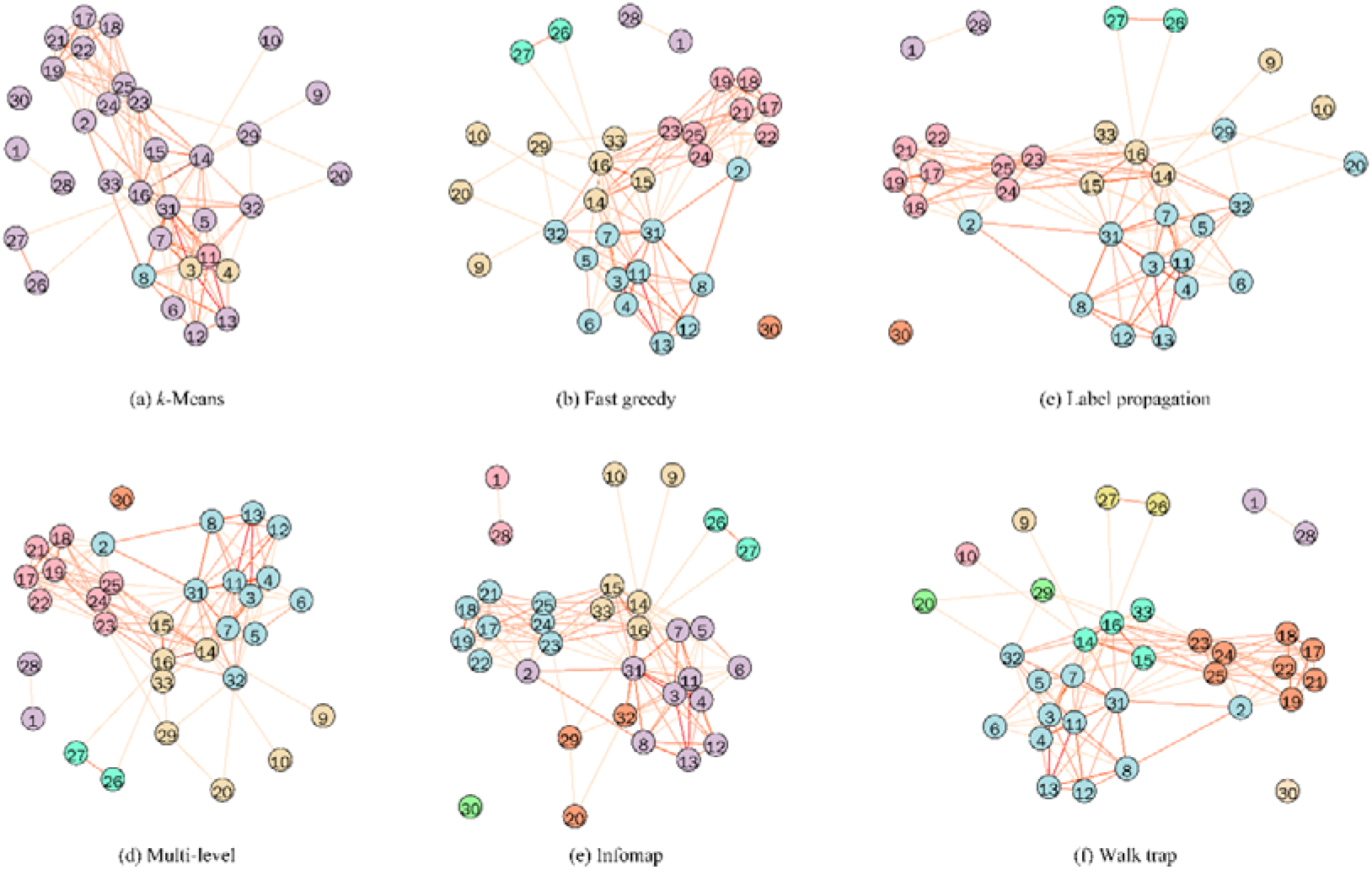
The label partition results that community detection algorithms and benchmark *k*-means method applied in the label space of the nuclear receptor data set.

On the other hand, we discuss the pathway and classification of three communities through the KEGG database, and the details shown in Table 5. The (1, 28) and (26, 27) are communities obtained by all six algorithms, and (20, 29, 32) are only available in IMA. For each of the first two communities, the two vertices belong to the same classification and pathway. The three vertices in the third community have similarities and differences. Therefore, we can think that the label clustering obtained by the community detection algorithm has a certain biological interpretation significance. This also confirms the classical assumption that similar targets tend to combine similar drugs.

**Table 5.**
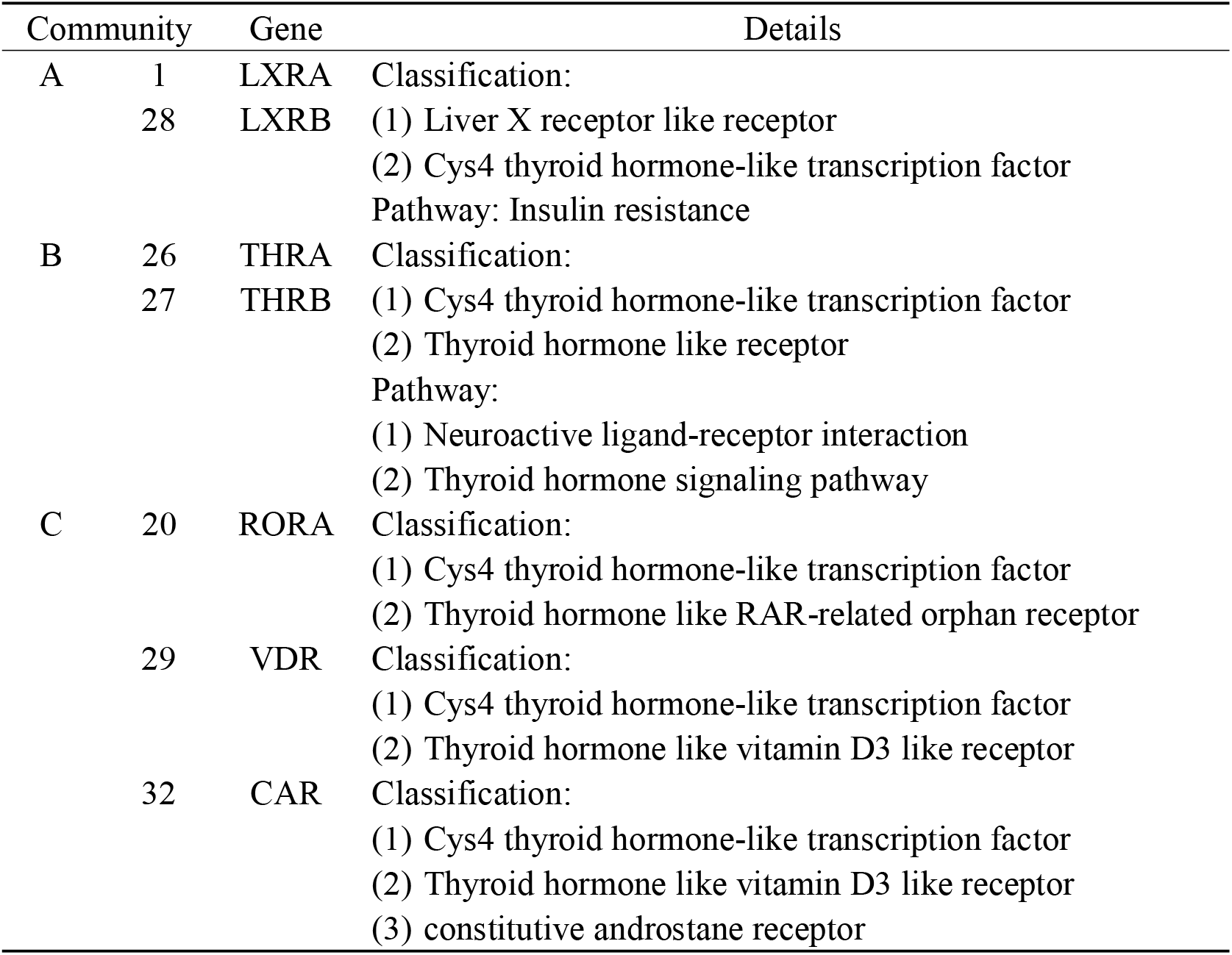
The details for three communities A: (1, 28), B: (26, 27), and C: (20, 29, 32). The numbers represent the nodes in Figure 7.

### 3.4. The DTI-MLCD and benchmark methods in Yamanishi_08 data sets

We compare the proposed method against three state-of-the-art methods for DTIs prediction. NetLapRLS [65], BLM-NII [116], and DDR [27]. NetLapRLS introduces the drug-target network information into the manifold Laplacian regularized the least square method which uses the concept of the bipartite local model. BLM-NII exploits bipartite local model with neighbor-based interaction profile inferring, which adds a preprocessing component to infer training data from neighbors’ interaction profiles. DDR executes the graph mining technique firstly to acquire the comprehensive feature vectors and then applies the random forest model by using different graph-based features extracted from the drug-target heterogeneous graph. Since these methods are proposed on the Yamanishi_08 data set, we perform the proposed DTI-MLCD method on this data set and compare it with other methods. All methods are carried out under the same experimental environment, such as SCV, random seeds, etc. And the results are obtained after fine-tuning. As reflected in Table 6, all the proposed methods in task T_D_ outperform the benchmark methods in terms of AUPR. For task T_T_ (Table 7), the proposed methods outperform benchmark methods in IC and E data sets while they are slightly inferior to BLM-NII in NR and GPCR. In order to comprehensively test the superiority of the method proposed in this study, we conduct the Friedman-Nemenyi test for all 8 methods. This hypothesis test is performed on both AUPR and AUC for completeness although AUPR is more informative than AUC in this study. And the result reveals that all the proposed methods are ranked before the three benchmark methods. Moreover, they are significantly better than DDR and NetLapRLS with a significance level of 0.05 and 0.1, respectively.

**Table 6.** The results of the proposed methods and three benchmark methods for task T_D_ (i.e. predicting new drugs). WTA, IMA, LPA, MLA, and FGA are community detection algorithms in the proposed DTI-MLCD method.

| Algorithm | AUC |  |  |  | AUPR |  |  |  |
| --- | --- | --- | --- | --- | --- | --- | --- | --- |
|  | NR | GPCR | IC | E | NR | GPCR | IC | E |
| FGA | 0.7829 | 0.8636 | 0.8220 | 0.8506 | 0.4990 | 0.4504 | 0.3887 | **0.4105** |
| IMA | 0.7830 | **0.8698** | 0.8223 | 0.8537 | 0.4992 | **0.4593** | 0.3857 | 0.4045 |
| LPA | 0.7785 | 0.8655 | 0.8197 | **0.8563** | **0.5079** | 0.4537 | **0.3924** | 0.4067 |
| MLA | 0.7829 | 0.8632 | **0.8237** | 0.8522 | 0.4990 | 0.4488 | 0.3885 | 0.4088 |
| WTA | 0.7828 | 0.8619 | 0.8219 | 0.8539 | 0.4989 | 0.4501 | 0.3860 | 0.4045 |
| BLM-NII | **0.8042** | 0.8496 | 0.8119 | 0.8204 | 0.4503 | 0.3415 | 0.3260 | 0.2690 |
| NetLapRLS | 0.7919 | 0.8281 | 0.7721 | 0.7933 | 0.4313 | 0.2456 | 0.2078 | 0.1287 |
| DDR | 0.6019 | 0.5678 | 0.4994 | 0.4768 | 0.2878 | 0.1907 | 0.1471 | 0.1336 |

**Table 7.** The results of the proposed methods and three benchmark methods for task T_T_ (i.e. predicting new targets). WTA, IMA, LPA, MLA, and FGA are community detection algorithms in the proposed DTI-MLCD method.

| Algorithm | AUC |  |  |  | AUPR |  |  |  |
| --- | --- | --- | --- | --- | --- | --- | --- | --- |
|  | NR | GPCR | IC | E | NR | GPCR | IC | E |
| FGA | 0.4961 | 0.7458 | 0.9104 | 0.9285 | 0.3472 | 0.2943 | 0.7047 | 0.7861 |
| IMA | 0.4929 | 0.7429 | **0.9114** | 0.9214 | 0.3457 | 0.2919 | 0.7027 | 0.7875 |
| LPA | 0.4925 | 0.7509 | 0.9105 | 0.9214 | 0.3398 | 0.2969 | 0.7082 | **0.7877** |
| MLA | 0.4998 | 0.7481 | 0.9098 | **0.9286** | 0.3487 | 0.2942 | **0.7093** | 0.7868 |
| WTA | 0.4923 | 0.7495 | 0.9103 | 0.9217 | 0.3460 | 0.3010 | 0.7046 | 0.7873 |
| BLM-NII | **0.5042** | **0.7777** | 0.9093 | 0.9193 | **0.3726** | **0.3078** | 0.7028 | 0.7570 |
| NetLapRLS | 0.4986 | 0.7425 | 0.9082 | 0.9161 | 0.2793 | 0.2515 | 0.6543 | 0.7064 |
| DDR | 0.4932 | 0.6290 | 0.5784 | 0.6965 | 0.2365 | 0.2288 | 0.3108 | 0.5026 |

### 3.5. Independent test

We conduct independent tests of the proposed DTI-MLCD method according to the data set before and after the update. First, build the independent test set. Drugs and their DTIs that do not exist in the Yamanishi_08 data set but exist in the updated data set will be used as independent test samples for task T_D_. Similarly, independent test samples of task T_T_ is constructed. Then, conduct independent tests on the model which trained on the Yamanishi_08 data set. The results are shown in Table 8 (task T_D_) and Table 9 (task T_T_).

**Table 8.** The results of independent tests on Yamanishi_08 data sets for task T_D_.

| Algorithm | AUC |  |  |  | AUPR |  |  |  |
| --- | --- | --- | --- | --- | --- | --- | --- | --- |
|  | NR | GPCR | IC | E | NR | GPCR | IC | E |
| FGA | 0.8174 | 0.8941 | 0.8238 | 0.8457 | 0.5331 | 0.3953 | 0.2795 | 0.1369 |
| IMA | 0.8172 | 0.9020 | 0.8262 | 0.8426 | 0.5331 | **0.4000** | 0.3012 | 0.1353 |
| LPA | 0.8157 | 0.9000 | 0.8257 | 0.8430 | **0.5334** | 0.3982 | **0.3013** | 0.1375 |
| MLA | 0.8174 | 0.8944 | 0.8246 | 0.8455 | 0.5331 | 0.3928 | 0.2776 | **0.1378** |
| WTA | 0.8174 | 0.8920 | 0.8230 | 0.8427 | 0.5331 | 0.3935 | 0.2890 | 0.1363 |

**Table 9.** The results of independent tests on Yamanishi_08 data sets for task T_T_.

| Algorithm | AUC |  |  |  | AUPR |  |  |  |
| --- | --- | --- | --- | --- | --- | --- | --- | --- |
|  | NR | GPCR | IC | E | NR | GPCR | IC | E |
| FGA | 0.8224 | 0.6130 | 0.7353 | 0.7348 | 0.3787 | 0.0076 | 0.2090 | **0.1077** |
| IMA | 0.8224 | 0.6135 | 0.7323 | 0.6834 | 0.3787 | 0.0075 | 0.2144 | 0.1057 |
| LPA | 0.8223 | 0.6107 | 0.7383 | 0.6809 | **0.3840** | 0.0076 | 0.2127 | 0.1048 |
| MLA | 0.8228 | 0.6255 | 0.7395 | 0.7339 | 0.3787 | **0.0080** | 0.2119 | 0.1071 |
| WTA | 0.8224 | 0.6080 | 0.7363 | 0.6814 | 0.3787 | 0.0074 | **0.2142** | 0.1052 |

## 4. Conclusion

This study updated the gold standard data set Yamanishi_08, and proposed DTI-MLCD for DTIs prediction, which is a new multi-label learning framework empowered by community detection. This framework has 5 effective models corresponding to five community detection algorithms to do label partitioning. This study conducted experiments on the gold standard data set before and after the update. On the original data set, the DTI-MLCD is compared with other benchmark methods, and its superiority is confirmed. In the updated data set, DTI-MLCD is superior to other classic machine learning algorithms. In addition, this study also constructed independent tests with new and old data sets. On the other hand, the results of the five community detection algorithms used in this framework are not significantly different. Moreover, they are superior to the benchmark *k*-means algorithm in performance and interpretability.

In the future, we will solve the problem of label imbalance and construct positive and negative samples in the form of semi-supervised learning to improve the performance of the framework in predicting DTIs.

